# Resource optimization in COVID-19 diagnosis

**DOI:** 10.1101/2020.06.25.172528

**Authors:** S.A Taniwaki, S.O.S Silva, N.F Santana-Clavijo, J. A Conselheiro, G.T Barone, A.A.R Menezes, E.S Pereira, P.E Brandão

**Affiliations:** Laboratory of Applied Molecular Biology and Serology, Department of Preventive Veterinary Medicine and Animal Health, School of Veterinary Medicine, University of São Paulo (USP), São Paulo – SP, Brazil; Laboratory of Diagnostics of Zoonosis and Vector-Borne Diseases, Zoonosis Surveillance Division, Health Surveillance Coordination, Municipal Health Secretariat, São Paulo – SP, Brazil

**Keywords:** SARS-CoV-2, COVID-19, RT-qPCR, cotton swabs, reaction volume

## Abstract

The emergence and rapid dissemination worldwide of a novel *Coronavirus* (SARS-CoV-2) results in decrease of swabs availability for clinical samples collection, as well as, reagents for RT-qPCR diagnostic kits considered a confirmatory test for COVID-19 infection. This scenario, showed the requirement of improve de diagnostic capacity, so the aim of this study were to verify the possibility of reducing the reaction volume of RT-qPCR and to test cotton swabs as alternative for sample collection. RT-qPCR volumes and RNA sample concentration were optimized without affecting the sensitivity of assays, using both probe-based and intercalation dyes methods. Although rayon swabs showed better performance, cotton swabs could be used as alternative type for clinical sample collection. COVID-19 laboratory diagnosis is important to isolate and restrict the dissemination of virus, so seek for alternatives to decrease the coast of assays improve the control of disease.

## 1. Introduction

On December 31, 2019, the World Health Organization (WHO) was notified about cases of severe pneumonia in Wuhan, China (WHO, 2020a). The etiological agent was identified as a coronavirus (*Coronaviridae: Betacoronavirus: Sarbecovirus: Severe acute respiratory syndrome-related coronavirus*) named as SARS-CoV-2 (GORBALENYA et al., 2020; LU et al., 2020a) and the disease denominated as COVID-19 (*Coronavirus disease* 2019) (WHO, 2020b). Confirmed cases are currently peaking in Latin America and, for instance, on June 13th 2020, more than 850,000 cases had been confirmed in Brazil (available in https://covid.saude.gov.br/).

The diagnostic test recommended for SARS-CoV-2 is a Real-Time Polymerase Chain Reaction (RT-qPCR), which can detect active infection by detection of viral genome. Collection of only nasopharyngeal samples (1 swab used for both nostrils) or combined with oropharyngeal (1 swab) sample, should be made with synthetic fibers swabs as more reliable due to a lesser interference with the downstream reactions (CDC, 2020a). Protocols with primers and probes designed at the Charité Virology Institute, Berlin (CORMAN et al., 2020) or at the Centers of Disease Control and Prevention (CDC,2020b; LU et al., 2020b) became one of the standard procedures for SARS-CoV-2 detection.

With the increasing demand for RT-qPCR tests for both patients and population screening, there is an ongoing shortage of both RT-qPCR reagents and rayon swabs; besides, the relatively high costs of these consumables for poor countries pose a barrier for a wider use of testing. This manuscript reports on the optimization of the Charité and the CDC RT-qPCR protocols for SARS-CoV-2 detection regarding concentration and volumes of reagents for both probe and intercalant agent-based platforms, as well as on the substitution of rayon swabs for cotton swabs for sample collection.

## 2. Materials and Methods and Results

### 2.1 Probe protocol: optimization of reaction volume and sample concentration

An inactivated SARS-CoV-2 isolate (kindly provided by Prof. Edison L. Durigon, Institute of Biomedical Sciences, University of São Paulo, Brazil) was used for total RNA extraction with QIAamp^®^ Viral RNA Mini kit (QIAGEN).

Total RNA concentrations of 10 and 20% for final reactions of 20, 15 and10 µL, were tested using AgPath-ID One Step RT-PCR kit (Applied Biosystems™) and StepOne™ Real-Time PCR Systems (Applied Biosystems™), with primers and probes for the RdRp and E genes (CORMAN et al., 2020). Amplification conditions were 48 °C/20 min (reverse transcription), 95 °C/10 min (for the activation of the DNA polymerase) followed by 45 cycles at 95 °C/ 10 seconds and 60 °C/ 30 seconds.

Though the different total RNA concentrations resulted in 1 Cq difference, no difference was found for the three final reaction volumes (Table 1).

**Table 1.** Cq mean for probe-based RT qPCR for the E and RdRp genes of SARS-CoV-2 with a range of RNA concentration and final reaction volumes.

| E gene |  |  | RdRp gene |  |  |
| --- | --- | --- | --- | --- | --- |
| Final volume<br>(μL) | RNA<br>(μL) | Cq mean | Final volume<br>(μL) | RNA<br>(μL) | Cq mean |
| 20 | 4 | 12,31 | 20 | 4 | 14,60 |
|  | 2 | 13,41 |  | 2 | - |
| 15 | 3 | 12,32 | 15 | 3 | 14,58 |
|  | 1,5 | 13,39 |  | 1,5 | 15,77 |
| 10 | 2 | 12,47 | 10 | 2 | 15,00 |
|  | 1 | 13,51 |  | 1 | 15,58 |

Performance of E and RdRp genes of SARS-CoV-2 RT-qPCRs, based on final reaction volume of 10 µL with 2 µL of RNA (Table 1), were verified with a relative standard curve built with 10^−2^ to 10^−8^ dilutions of positive RNA control. Respectively for E and RdRp genes, detection were observed until 10^−6^ (Cq mean= 33.71, E= 91.2% and r^2^= 0.998) and 10^−5^ dilutions (Cq mean= 31.98, E= 96.82% and r^2^= 0.998). This lower sensitivity for RdRp gene is possibly a consequence of a lower transcription of the corresponding ORF due to a 3-to-5’ attenuation (IRIGOYEN et al., 2016). So, the results showed that the final volume does not interfere with the sensitivity of this assay.

The kits GoTaq^®^ probe 1-Step RT-qPCR (Promega) and SuperScript III Platinum One-Step RT-PCR System (ThermoFisher Scientific) were also tested for the final reaction volumes of 15 and 10 µL, as per manufacturer’s instructions, in a 7500 Real-Time PCR Systems (Applied Biosystems™), and the same results were found when compared to the AgPath-ID One Step RT-PCR kit (Applied Biosystems™).

For the CDC protocol (LU et al., 2020b), 2019-nCoV CDC-qualified Probe and Primer for SARS-CoV-2 kit (Biosearch(tm) Technologies), which includes primers and probes for SARS-CoV-2 N gene (N1 and N2) and Human RNase P (endogen control), were tested with final reaction volume of 20 and 10 µL with 25% concentration of total RNA, using AgPath-ID One Step RT-PCR (Applied Biosystems™) and StepOne™ Real-Time PCR Systems (Applied Biosystems™). Same conditions and relative standard curve (10^−2^ to 10^−^8) were used with E gene reaction (CORMAN et al., 2020) to possibly the comparison between assays performances.

All reactions resulted in adequate efficiencies and r^2^ values (Table 2) and detected the targets up to the 10^−8^ dilution (Table 2), though the N2 reaction resulted in a more intense fluorescence for the higher dilutions, allowing an easier visualization of amplification. The difference between the sensitivities of this last test with the previously described for the Charité RdRp and E genes protocol is possibly due to different positive control aliquots used.

**Table 2.** Performance for probe-based RT-qPCR assays for N (N1 and N2) and E genes of SARS-CoV-2.

| Dilution | Cq mean |  |  |  |  |
| --- | --- | --- | --- | --- | --- |
|  | E | N1 |  | N2 |  |
|  | 10 µL | 10 µL | 20 µL | 10 µL | 20 µL |
| 10<sup>-2</sup> | 18.65 | 18.59 | 18.49 | 18.90 | 18.35 |
| 10<sup>-3</sup> | 21.71 | 21.82 | 21.87 | 22.35 | 21.83 |
| 10<sup>-4</sup> | 25.10 | 24.84 | 25.15 | 25.34 | 25.27 |
| 10<sup>-5</sup> | 28.12 | 27.92 | 28.01 | 28.52 | 28.33 |
| 10<sup>-6</sup> | 31.32 | 31.30 | 31.32 | 32.35 | 31.66 |
| 10<sup>-7</sup> | 34.40 | 34.03 | 34.31 | 35.09 | 35.18 |
| 10<sup>-8</sup> | 37.79 | 36.96 | 38.39 | 38.26 | 38.38 |
| **Efficiency (%)** | 106.3 | 111.0 | 102.7 | 103.3 | 99.7 |
| **r<sup>2</sup>** | 0.993 | 0.999 | 0.998 | 0.998 | 0.999 |

The limit of detection (LOD) was calculated for each assay using 14 replicates of each dilution (1×10^−7^, 2×10^−8^ and 1×10^−8^). The N2 reaction showed slight better results with LOD of 2×10^−8^ dilution, while E and N1 assays had LOD of 1×10^−7^ dilution (Table 3).

**Table 3.** Limit of detection for probe-based RT-qPCR assays for N (N1 and N2) and E genes of SARS-CoV-2.

|  | 1x10<sup>-8</sup> |  |  | 2x10<sup>-8</sup> |  |  | 1x10<sup>-7</sup> |  |  |
| --- | --- | --- | --- | --- | --- | --- | --- | --- | --- |
|  | E | N1 | N2 | E | N1 | N2 | E | N1 | N2 |
| Positive | 5 | 4 | 9 | 6 | 6 | 11 | 14 | 11 | 13 |
| percentage | 35,71 | 28,57 | 64,29 | 42,9 | 42,9 | 78,6 | 100,0 | 78,6 | 92,9 |
| Ct mean | 39,06 | 36,97 | 37,66 | 38,81 | 37,04 | 37,57 | 36,37 | 36,03 | 36,19 |
| lower Ct | 36,74 | 36,16 | 36,35 | 37,19 | 36,01 | 35,45 | 34,12 | 34,96 | 35,03 |
| higher Ct | 41,53 | 37,90 | 38,20 | 40,64 | 37,31 | 39,21 | 38,21 | 38,60 | 38,07 |
| SD | 1,94 | 0,90 | 0,66 | 1,46 | 0,51 | 1,10 | 1,09 | 1,13 | 0,98 |

### 2.2 Intercalating dyes: optimization of primers concentration

For the intercalating dyes system, primers targeting human RNAse P, as per the CDC protocol (LU et al., 2020b), were used at 50, 100, 150, 200 and 250 nM each. For E and RdRp genes from the Charité protocol (CORMAN et al., 2020), primers were tested at 100 and 200 nM and 100, 200, 300 and 400 nM, respectively.

Reactions were carried out with *Power* SYBR^®^ Green RNA to Ct 1-Step kit (Applied Biosystems™) using a StepOne™ and 7500 Real-Time PCR Systems (Applied Biosystems™) equipment, using 3 or 2 µL of RNA for 15 and 10 µL final volume reactions, respectively. Cycling conditions were 48 °C/20 min (reverse transcription), 95 °C/10 min (for the activation of the DNA polymerase) followed by 45 cycles at 95 °C/ 10 seconds and 60 °C/ 30 seconds and melt curve analyses (95 °C/15 secs, 60 °C/15 secs and 95 °C/15 secs).

For the RNAse P, the 100 nM primers reaction was found as more efficient (E=101.8%; r^2^=0.991), with a specific peak at 81 °C in the melt curve; at higher primers concentrations, primers dimers were detected in melt curve analysis and at 50 nM there was a loss in sensitivity.

E gene reaction was fond as sensitive as the probe-based reaction for this target, being able to detect up to 10^−6^ dilution of positive RNA control with primers at both 100 and 200 nM, with a unique melting peak at 77 °C. At 200 nM primers concentration, though, a higher efficiency was found (E= 99.3 % and r^2^=0.998), as well as an earlier target detection for the 10^−6^ dilution (Cq mean= 32.53 and 30.07, respectively, for 100 and 200 nM).

The RdRp reaction showed insufficient performance, as, with primers at 200 nM there was a loss in sensitivity, while, at 400 nM the sensitivity was equal to the probe-based reaction (detection up to 10^−5^ dilution), but efficiency could not be evaluated due to primer dimers formation visualized in melt curve analysis.

### 2.3 Comparison between cotton and rayon swabs

An inactivated SARS-CoV-2 isolate was 2-fold diluted in sterile saline from 2^-1^ to 2^-9^ in 100 µL, in duplicate. For each respective dilution, one rayon (Inlab Diagnóstica, Brazil) and one cotton (Labor, Brazil) swab, both with plastic stalks, were dipped into the tube (one swab/tube), being all the content absorbed by the swabs. Undiluted SARS-CoV-2 (2^0^) was also included, as well as sterile saline (negative control).

Next, each swab was immersed in a conical tube containing 3 mL sterile saline and kept at 2-8 °C for 72 hours, in order to mirror the clinical flow of clinical samples. All swabs were finally submitted to RNA extraction and tested with the probe-based E gene RT-qPCR protocol.

Rayon swabs showed more consistency results, with detection through all dilutions from 2^0^ to 2^-9^ linearly (Table 4), while cotton swabs failed to detect in two dilutions (2^-5^ and 2^-6^). CDC does not recommend the use of calcium arginate swabs and swabs with wooden shafts, because these material types may contain PCR inhibitors, and indicate only use of synthetic fibers swabs for collection of clinical samples (CDC, 2020a). However, some study showed good performance of cotton swabs for detection of Human norovirus and rhinovirus by RT-qPCR (WARIS et al., 2013; LEE et al., 2018). Both cotton and rayon swabs have the same chemical structure, with O-H groups which form hydrogen bonds with nucleic acids, and showed optimal absorption capacity (above 100 µL), but low extraction and recovery efficiency (BRUIJNS, 2018). Although the amplification have been failed in some dilutions, in lower and higher dilution detection was possible, thus cotton swabs could be an alternative to rayon swabs for clinical sample collection.

**Table 4.**
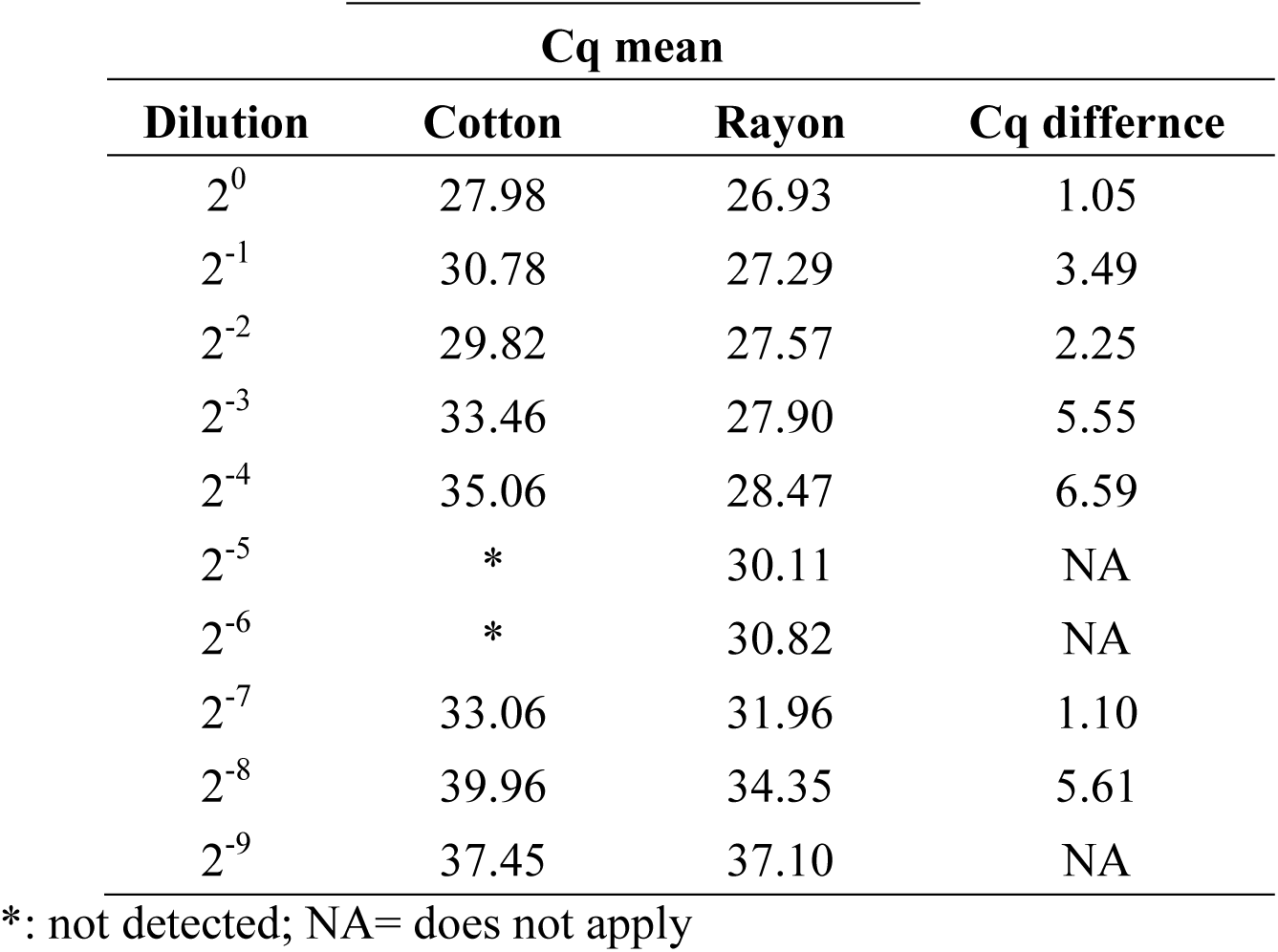
Probe-based RT-qPCR to the E gene of serial dilutions of SARS-CoV-2 sampled with cotton and rayon swabs.

| Dilution | Cq mean |  | Cq difference |
| --- | --- | --- | --- |
|  | Cotton | Rayon |  |
| 2<sup>0</sup> | 27.98 | 26.93 | 1.05 |
| 2<sup>-1</sup> | 30.78 | 27.29 | 3.49 |
| 2<sup>-2</sup> | 29.82 | 27.57 | 2.25 |
| 2<sup>-3</sup> | 33.46 | 27.90 | 5.55 |
| 2<sup>-4</sup> | 35.06 | 28.47 | 6.59 |
| 2<sup>-5</sup> | * | 30.11 | NA |
| 2<sup>-6</sup> | * | 30.82 | NA |
| 2<sup>-7</sup> | 33.06 | 31.96 | 1.10 |
| 2<sup>-8</sup> | 39.96 | 34.35 | 5.61 |
| 2<sup>-9</sup> | 37.45 | 37.10 | NA |

\*: not detected; NA= does not apply

## 3. Conclusions

The consequences of the ongoing shortage of rayon swabs and RT-qPCR reagents on the sampling and testing of COVID-19 suspected persons will surely have a negative impact on the control of the disease due to a lower sampling and a higher sub notification rate. Here, we demonstrated that the decrease of reaction volume not interferes in sensitivity of assays, allowing the diagnostic capacity increase, and a choice of the type of swab must be based in the local availability and the urgency of the sampling and testing.

## 4. Consolidated final protocols

Once no loss of sensitivity was demonstrated using 10 µL final volume reactions, the following protocols are suggested for probe-based and intercalating dyes RT-qPCRs for detection of SARS-CoV-2.

Amplification conditions were optimized for both probe and intercalating dye reactions and for different kits and primers sets.

| Stage | Step | Temperature | Time |
| --- | --- | --- | --- |
| Holding | Reverse transcription | 48 °C | 20 min |
| Holding | Polymerase activation | 95 °C | 10 min |
| Cycling<br>45x | Denaturing | 95 °C | 10 s |
|  | Annealing/Extension | 60 °C | 30 s |
| Melting<br>curve* | Denaturing | 95 °C | 15 s |
|  | Annealing | 60 °C | 15 s |
|  | Denaturing | 95 °C | 15 s |

\*applies only for the intercalating-dye based reactions

### For probes-based reactions

| SARS-CoV-2 E gene |  |  |
| --- | --- | --- |
| Reagent | Final concentration | Volume/sample (μL) |
| AgPath-ID Buffer (2x) | 1x | 5.00 |
| Primer E_Sarbeco_F1 (10 μM) | 400 nM | 0.40 |
| Primer E_Sarbeco_R2 (10 μM) | 400 nM | 0.40 |
| Probe E_Sarbeco_P1 (10 μM) | 200 nM | 0.20 |
| RT mix (25x) | 1x | 0.40 |
| H2O | to 10 μL | 1.10 |
| Total |  | 7.50 |
| RNA |  | 2.50 |

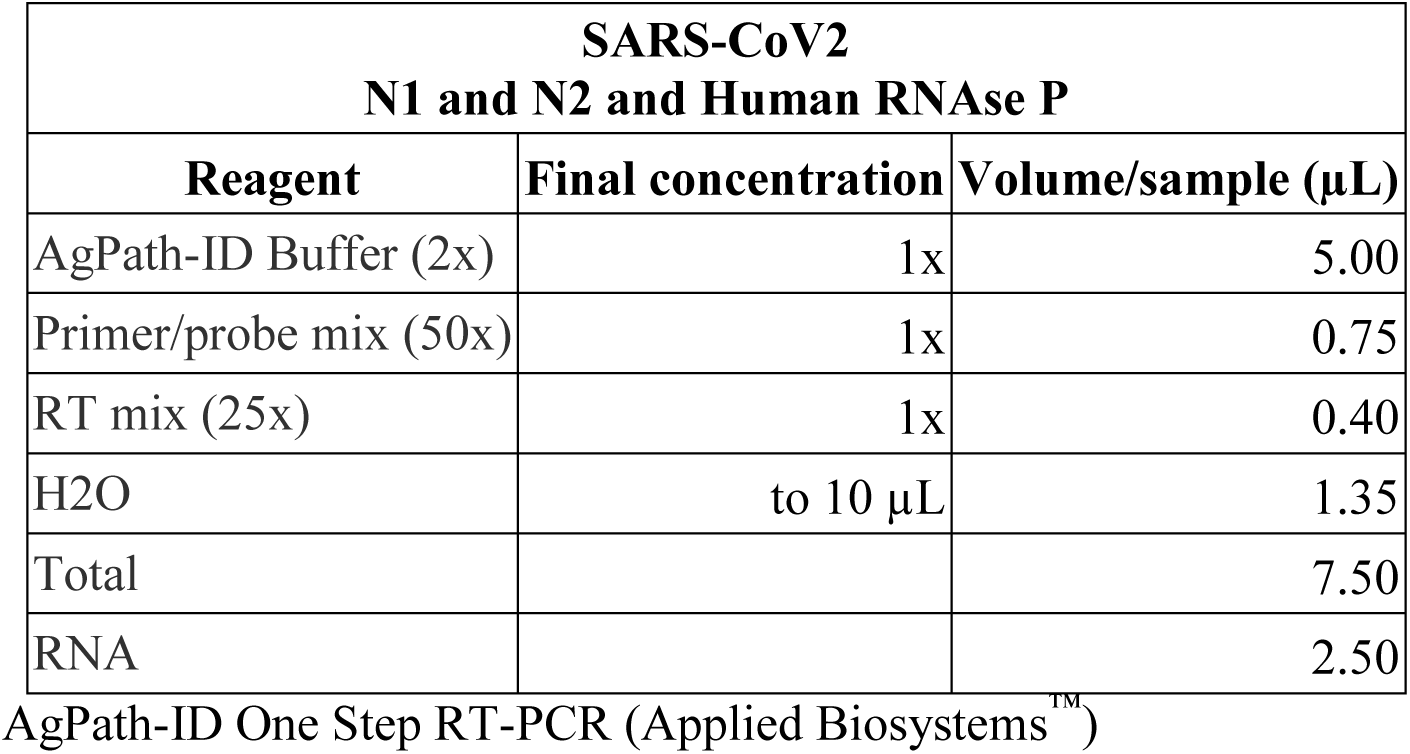

| **SARS-CoV2**<br>**N1 and N2 and Human RNase P** |  |  |
| --- | --- | --- |
| **Reagent** | **Final concentration** | **Volume/sample (μL)** |
| AgPath-ID Buffer (2x) | 1x | 5.00 |
| Primer/probe mix (50x) | 1x | 0.75 |
| RT mix (25x) | 1x | 0.40 |
| H2O | to 10 μL | 1.35 |
| Total |  | 7.50 |
| RNA |  | 2.50 |

### For intercalating dyes

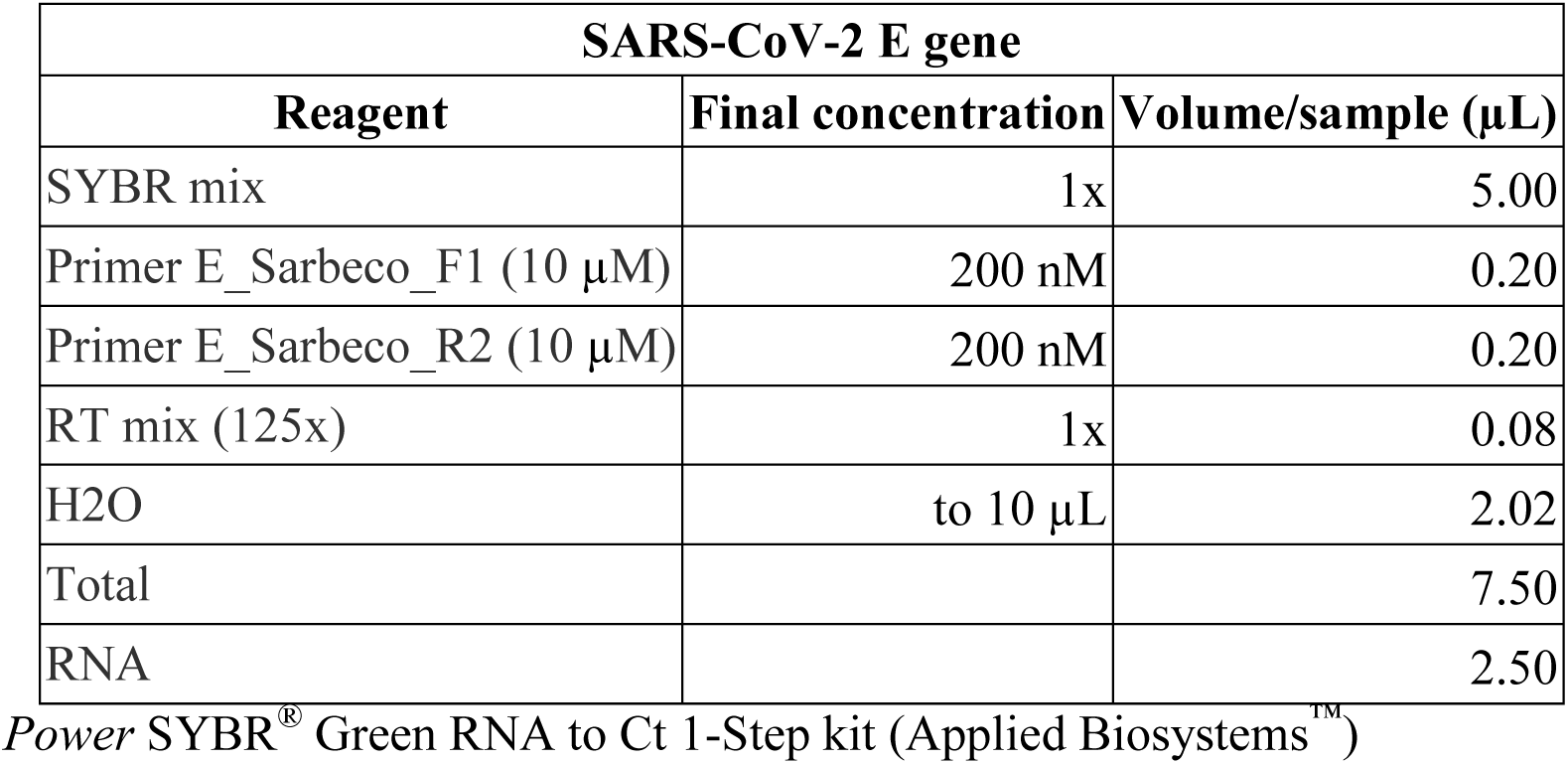

AgPath-ID One Step RT-PCR (Applied Biosystems™) - For intercalating dyes
| **SARS-CoV-2 E gene** |  |  |
| --- | --- | --- |
| **Reagent** | **Final concentration** | **Volume/sample (μL)** |
| SYBR mix | 1x | 5.00 |
| Primer E_Sarbeco_F1 (10 μM) | 200 nM | 0.20 |
| Primer E_Sarbeco_R2 (10 μM) | 200 nM | 0.20 |
| RT mix (125x) | 1x | 0.08 |
| H2O | to 10 μL | 2.02 |
| Total |  | 7.50 |
| RNA |  | 2.50 |

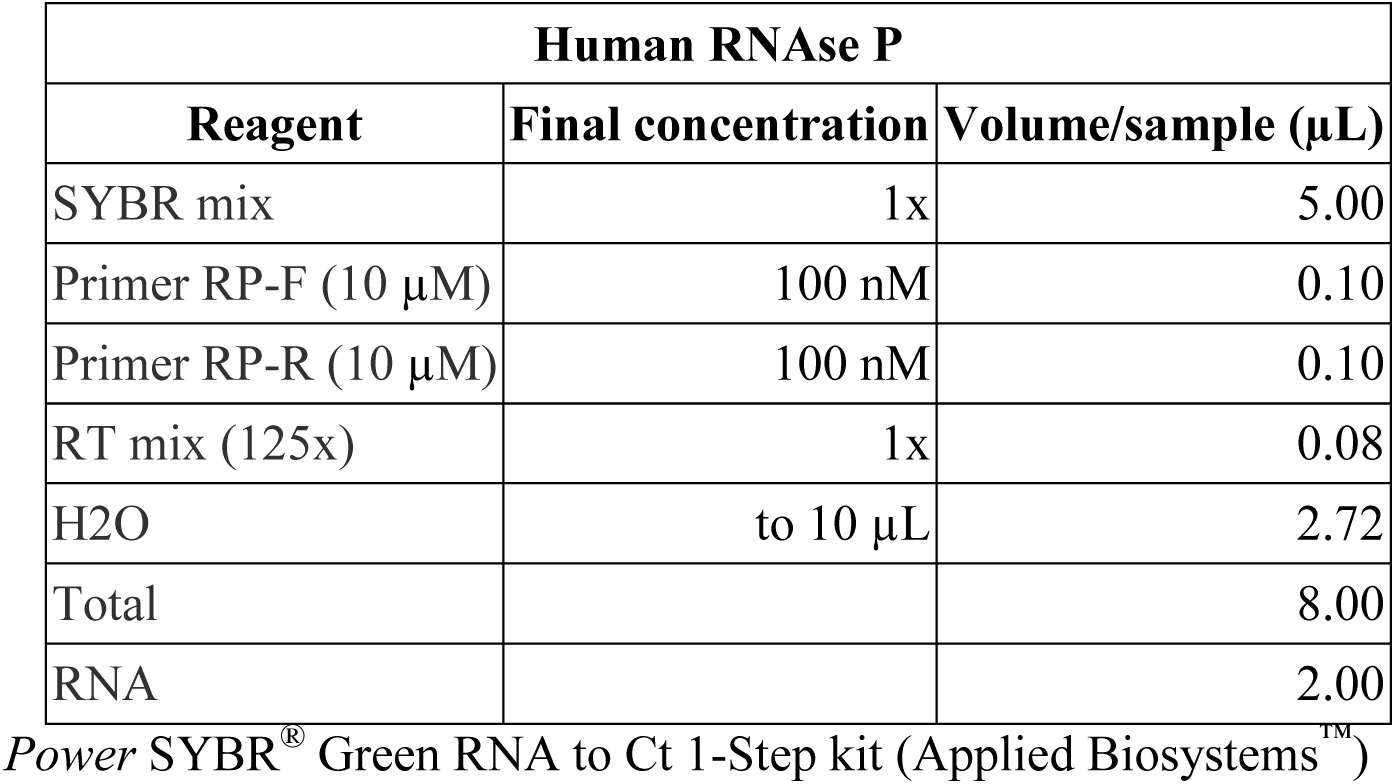

## 5. Acknowledgments

This work was funded by FAPESP grant number 2017/24907-6, CNPq (Brazilian National Board for Scientific and Technological Development grant number 307291/2017-0 and CAPES (Coordenação de Aperfeiçoamento de Pessoal de Nível Superior, Brasil - Finance Code 001). Authors were grateful to Prof. Edison L. Durigon and his team of Laboratory of Clinical and Molecular Virology, Institute of Biomedical Sciences, University of São Paulo, Brazil, for provided the inactivated SARS-CoV-2 isolate. And we thank Dr. Caroline Cotrim Aires from Laboratory of Diagnostics of Zoonosis and Vector-Borne Diseases, and Dr. Solange Maria de Saboia e Silva from Health Surveillance Coordination for the institutional support and incentive to the development of the study.

## Notes

### Competing Interest Statement

The authors have declared no competing interest.

